# Expression Profiling, Downstream Signaling and Subunit Interactions of GPA2/GPB5 in the Adult Mosquito *Aedes aegypti*

**DOI:** 10.1101/694653

**Authors:** David A. Rocco, Jean-Paul V. Paluzzi

## Abstract

GPA2/GPB5 and its receptor constitute a glycoprotein hormone-signalling system native to the genomes of most vertebrate and invertebrate organisms, including humans and mosquitoes. Unlike the well-studied gonadotropins and thyrotropin, the exact function of GPA2/GPB5 is unclear, and whether it elicits its functions as heterodimers, homodimers or as independent monomers remains unknown. Here, the glycoprotein hormone signalling system was investigated in adult mosquitoes, where GPA2 and GPB5 subunit transcripts co-localized to bilateral pairs of neuroendocrine cells within the first five abdominal ganglia of the central nervous system. Unlike human GPA2/GPB5 that demonstrated strong heterodimerization between subunits, the GPA2/GPB5 subunits in *A. aegypti* lacked evidence of heterodimerization when heterologously expressed. Interestingly, cross-linking analysis to determine subunit interactions revealed *A. aegypti* and *H. sapiens* GPA2 and GPB5 subunits form homodimers, and treatments with independent subunits did not activate *A. aegypti* LGR1 or *H. sapiens* TSH receptor, respectively. Since mosquito GPA2/GPB5 heterodimers were not evident by heterologous expression of independent subunits, a tethered construct was generated for expression of the subunits as a single polypeptide chain to improve heterodimer formation. Our findings revealed *A. aegypti* LGR1 elicited constitutive activity that elevated levels of cAMP as determined by increased cAMP-dependent luminescence. However, upon treatment with recombinant tethered GPA2/GPB5 heterodimers, an inhibitory G protein (Gi) signalling cascade is initiated and forskolin-induced cAMP production is inhibited. These results provide evidence towards the functional deorphanization of LGR1 and, moreover, further support the notion that GPA2/GPB5 heterodimerization is a requirement for glycoprotein hormone receptor activation.

## Introduction

Members of the cystine knot growth factor (CKGF) superfamily, which are characterized with a CKGF domain as their primary structural feature, include (i) the glycoprotein hormones, (ii) the invertebrate bursicon hormone, (iii) the transforming growth factor beta (TGFβ) family, (iv) the bone morphogenetic protein (BMP) antagonist family, (v) the platelet-derived growth factor (PDGF) family and (vi) the nerve growth factor (NGF) family. Of the members of the CKGF superfamily, the glycoprotein hormones are of fundamental importance in the regulation of both vertebrate and invertebrate physiology.

In vertebrates, members of the glycoprotein hormone family include follicle-stimulating hormone (FSH), luteinizing hormone (LH), thyroid-stimulating hormone (TSH) as well as chorionic gonadotropin (CG), which are implicated in governing several aspects of physiology including reproduction, energy metabolism along with growth and development. Structurally, these hormones are formed by the heterodimerization of two cystine-knot glycoprotein subunits, an α subunit that is structurally identical for each hormone (GPA1), and a hormone-specific β subunit (GPB1-4) ^1, 2^.

Two novel glycoprotein hormone subunits were identified in the human genome, glycoprotein α2 (GPA2) and glycoprotein β5 (GPB5), and were found to heterodimerize (GPA2/GPB5) and act on the same receptor as TSH. As a result, GPA2/GPB5 was coined the name thyrostimulin to differentiate it from TSH in vertebrates ^3, 4^. Unlike other glycoprotein hormones which are restricted to the vertebrate lineage, homologous genes encoding GPA2/GPB5 subunits exist in all bilaterian organisms, where its function appears to be pleiotropic ^3–5^. In vertebrates, the function of GPA2/GPB5 has been implicated, or at least suggested, to be involved in reproduction ^6^, thyroxine production ^3, 7^, skeletal development ^8^, immunoregulation ^9, 10^ and the proliferation of ovarian cancer cell lines ^11^. For invertebrate species, GPA2/GPB5 has been implicated or demonstrated to function in development ^12–15^, hydromineral balance ^13–18^ as well as in reproduction ^18, 19^.

Non-covalent interactions and heterodimerization between the subunits forming FSH, LH, TSH and CG is required for their respective biological functions ^1, 20^. However, whether heterodimerization is required for GPA2/GPB5 to activate its receptor and exert its physiological role in vertebrates and invertebrates is debated. For FSH, LH, TSH and CG, the subunits are co-expressed in the same cells and each hormone is released into circulation as heterodimers ^21, 22^. On the other hand, GPA2 and GPB5 subunit expression profiles, both for vertebrate and invertebrate organisms, do not always occur in the same cells, and GPA2 is often expressed more widely and abundantly than GPB5 in some tissues ^7, 12, 23–25^. Additionally, unlike the beta subunits of the classic glycoprotein hormones, the structure of the GPB5 subunit lacks an extra pair of cysteine residues that form an additional disulfide linkage, referred to as the ‘seatbelt’, which strengthens and stabilizes its heterodimeric association with GPA2 ^26^.

Relative to other G protein-coupled receptors (GPCRs), members of the leucine-rich repeat-containing G protein-coupled receptor (LGR) family are often characterized with a large extracellular amino terminal domain responsible for the selective binding of their large hormone ligands ^27^. Following its initial genomic and molecular characterization ^28^, an invertebrate receptor for GPA2/GPB5, called LGR1, was functionally deorphanized in the fruit fly *Drosophila melanogaster*, where GPA2/GPB5 heterodimers were found to activate LGR1 and increased intracellular levels of cyclic AMP (cAMP) ^29^. Interestingly, stimulatory G protein (Gs) coupling and signalling to elevate cAMP was also shown with GPA2/GPB5-TSH receptor activation in humans ^3, 6^.

In the mosquito *Aedes aegypti*, genes encoding for GPA2, GPB5 and LGR1 were identified and shown to be expressed in all developmental life stages, with expression enriched in adults compared to juvenile stages ^14^. In adults, LGR1 was found localized to epithelia throughout the gut where GPA2/GPB5 could regulate feeding-related processes and hydromineral balance ^18^. Notably, LGR1 transcript expression was also observed in the reproductive organs of males and females ^19^. In adult male mosquitoes, knockdown of LGR1 expression led to abnormal spermatogenesis with spermatozoa displaying malformations such as shortened flagella and consequently, LGR1-knockdown males had 60% less spermatozoa as well as significantly reduced fecundity relative to control mosquitoes ^19^.

With an interest in better understanding GPA2/GPB5 signalling in *A. aegypti* mosquitoes, our work herein set out to characterize the tissue-specific and cellular distribution expression profile of GPA2/GPB5 in mosquitoes. As well, we sought to determine GPA2/GPB5 interactions and demonstrate GPA2/GPB5-LGR1 functional coupling using a heterologous system. Combining various molecular techniques, we demonstrate GPA2/GPB5 cellular co-expression in the central nervous system of adult mosquitoes, and that heterodimers are indeed required to activate LGR1, which exhibits ligand-dependent G protein-coupling activity. Moreover, these results provide evidence for human and mosquito GPA2 and GPB5 homodimers, which were incapable of activating TSH receptor or LGR1, respectively. Overall, these findings appreciably advance our understanding of GPA2/GPB5 signalling in mosquitoes and provide novel directions to uncover the functions of homologous systems in other organisms.

## Results

### *A. aegypti* GPA2 and GPB5 subunit expression localization

To determine the distribution of GPA2 and GPB5 subunit expression in the central nervous system and peripheral tissues, adult mosquito organs were analyzed using RT-qPCR. GPA2 and GPB5 subunit transcript was detected in the central nervous system of adult male and female mosquitoes, with significantly enriched expression in the abdominal ganglia relative to other nervous and peripheral tissues (Fig. 1a, b). Fluorescence *in situ* hybridization was employed to localize cell-specific expression of the GPA2 and GPB5 transcripts in the abdominal ganglia. GPA2 and GPB5 anti-sense RNA probes identified two bilateral pairs of neuroendocrine cells (Fig. 1c, d) within each of the first five abdominal ganglia in male and female mosquitoes whereas the control sense probes did not detect cells in the nervous system (Fig. 1e, f). Within each of these first five abdominal ganglia, GPA2 transcript localized to similar laterally-positioned cells as the GPB5 transcript (Fig. 1g, h). To determine if cells expressing GPA2 transcript were the same cells expressing GPB5 transcript, abdominal ganglia were simultaneously treated with both GPA2 and GPB5 anti-sense RNA probes. Using this dual probe approach, results confirmed the detection of only two bilateral pairs of cells (Fig. 1i, j) that displayed a greater staining intensity compared to the intensity of cells detected when using either the GPA2 or GPB5 anti-sense probe alone (Fig. 1c, d).

**Fig. 1.**
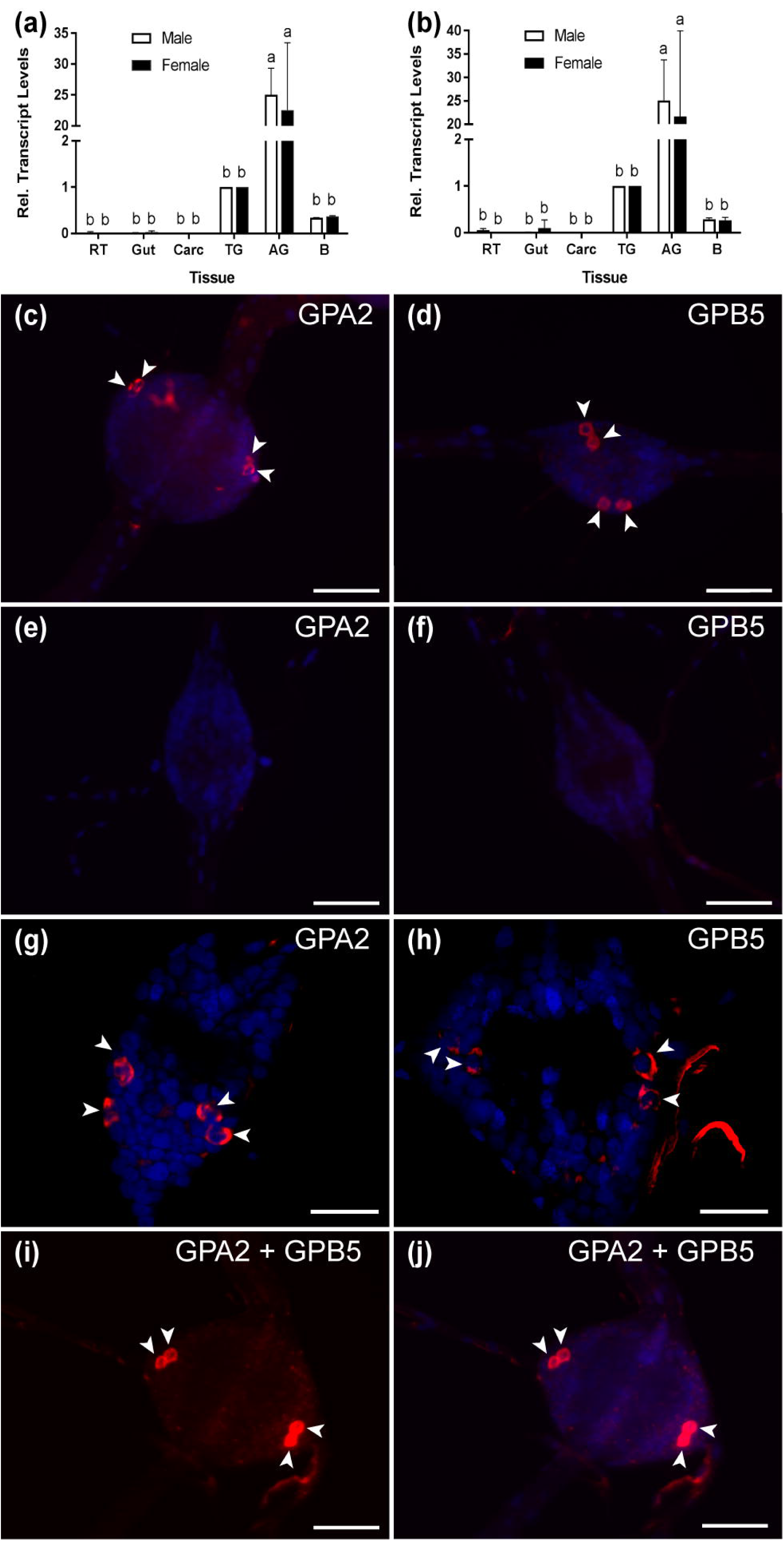
GPA2/GPB5 subunit transcript expression and localization in adult *A. aegypti*. (a, b) RT-qPCR examining GPA2 (a) and GPB5 (b) transcript expression in the central nervous system of adult mosquitoes, with significant enrichment in the abdominal ganglia (AG). Subunit transcript abundance is shown relative to their expression in the thoracic ganglia (TG). Mean ± SEM of three biological replicates. Columns denoted with different letters are significantly different from one another. Multiple comparisons two-way ANOVA test with Tukey’s multiple comparisons (P<0.05) to determine sex- and tissue-specific differences. Fluorescence *in situ* hybridization anti-sense (c, d, g-j) and sense (e, f) probes to determine GPA2 and GPB5 transcript localization (GPA2 and/or GPB5 transcript, red; nuclei, blue). Unlike sense probe controls (e, f), two bilateral pairs of cells (arrowheads) were detected with GPA2 (c, g) and GPB5 (d, h) anti-sense probes in the first five abdominal ganglia. Co-localization of the GPA2 (g) and GPB5 (h) transcript was verified by treating abdominal ganglia dually with a combination of GPA2 and GPB5 anti-sense probe (i, j) that revealed two, intensely-stained bilateral pairs of cells. In (c-f) and (i-j), microscope settings were kept identical when acquiring images of control and experimental ganglia. Scale bars are 50 µm in (c-f), (i-j) and 40 µm in (g) and (h).

Using a custom antibody targeting *A. aegypti* GPB5, we next sought to immunolocalize GPB5 protein in the abdominal ganglia. GPB5 immunoreactivity localized to two bilateral pairs of cells (Fig. 2a) within the first five ganglia, which were in similar positions to cells expressing GPA2 and GPB5 transcript (Fig. 1c, d, g-j). In some preparations, three bilateral pairs of cells immunolocalized to the first five abdominal ganglia of the ventral nerve cord (Fig. 2b), however no cells were ever detected in the sixth terminal ganglion (Fig. 2c). Along the lateral sides of each of the first five abdominal ganglia, GPB5 immunoreactive processes were observed to interconnect and closely associate as a tract of axons emanating through the lateral nerve (Fig. 2b). Control treatments with GPB5 antibody preabsorbed with the GPB5 immunogenic antigen, failed to detect any cells in ganglia (Fig. 2d)

**Fig. 2.**
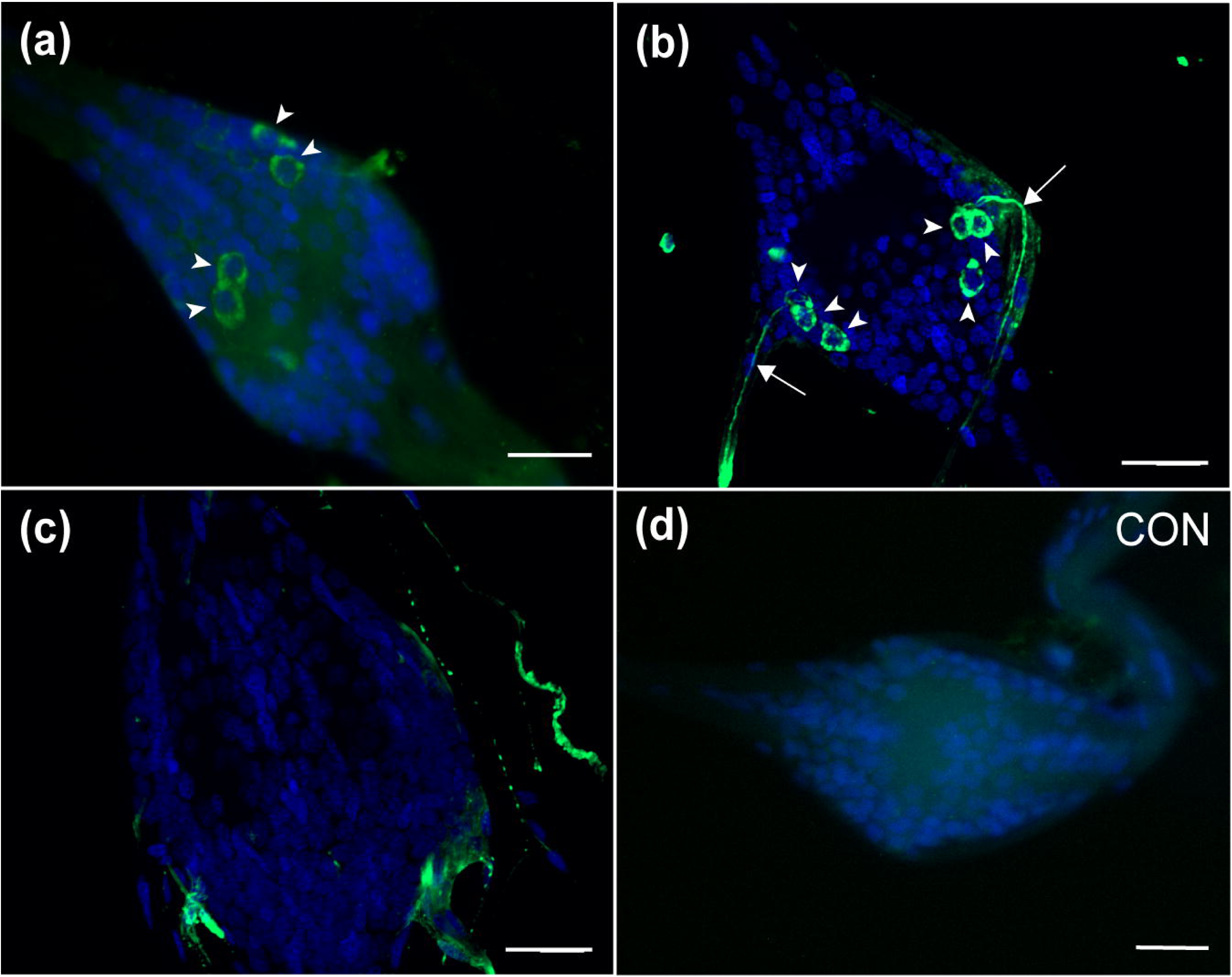
Immunolocalization of GPB5 subunit expression in the abdominal ganglia of adult *A. aegypti*. Experimental treatments demonstrated GPB5 immunoreactivity (green) in two (a), and in some mosquitoes, three (b) bilateral pairs of neuroendocrine cells (arrowheads) within the first five abdominal ganglia of the ventral nerve cord (DAPI, blue). Optical sections of ganglia revealed axonal projections emanate from GPB5-immunoreactive cells through the lateral nerves (arrows). No cells were detected in the sixth, terminal ganglion (c) and in control treatments (CON) where GPB5 antibody was preabsorbed with GPB5 synthetic antigen (d). In (a, d) or (b-c), microscope settings were kept identical when acquiring images of control and experimental ganglia. Scale bars are 25 µm in (a-d) and 20 µm in (b-c).

### Cross-linking analyses to determine *A. aegypti* GPA2/GPB5 dimerization patterns

*A. aegypti* GPA2 and GPB5 subunit protein interactions were studied using western blots of recombinant proteins from HEK 293T cells expressing each subunit independently, or co-expressing both subunits in the same cells using a dual promoter plasmid. Under control conditions, GPA2 protein is represented as two bands at 16 kDa and 13 kDa, which correspond to the glycosylated and non-glycosylated forms of *A. aegypti* GPA2, respectively (Fig. 3a). Following deglycosylation with PNGase, the higher molecular weight band of GPA2 is eliminated, and the non-glycosylated lower molecular weight band intensifies (Fig. 3a). Interestingly, when the GPA2 subunit was tested to examine potential homodimerization, an additional strong band at ∼32 kDa was detected, which migrates to ∼30 kDa when cross-linked protein samples were deglycosylated using PNGase (Fig. 3a). Under control conditions, GPB5 protein is represented as a band size at 24 kDa and the migration pattern is not affected by PNGase treatment (Fig. 3b). Upon treatment with DSS, a faint second band appears at 48 kDa, which does not change in molecular weight following treatment with PNGase (Fig. 3b). Three independent band sizes at 24 kDa (GPB5), 16 kDa (glycosylated GPA2) and 13 kDa (non-glycosylated GPA2) were detected in lanes loaded with protein isolated from HEK 293T cells co-expressing GPA2 and GPB5 subunits (Fig. 3c). After removing N-linked sugars, the 24 kDa band is not affected but the 16 kDa band disappears and 13 kDa band intensifies (Fig. 3c), as observed when assessing the GPA2 subunit independently (Fig. 3a). Cross-linked samples show the addition of two higher molecular weight bands at ∼48 kDa and ∼32 kDa, the latter of which migrates lower to ∼30 kDa when subjected to PNGase treatment (Fig. 3c). Given the resolution and intensity of the detected bands, molecular weight bands at ∼48 kDa and ∼32 kDa could indicate GPA2/ GPB5 heterodimeric interactions (13 or 16 kDa + 24 kDa = 37-40 kDa). Alternatively, these bands could reflect GPA2 (13 or 16 kDa + 13 or 16 kDa = 26-32 kDa) and GPB5 (24 + 24kDa = 48 kDa) homodimers. As a result, to clarify whether *A. aegypti* GPA2 and GPB5 subunits are heterodimeric partners, additional experiments were performed.

**Fig. 3.**
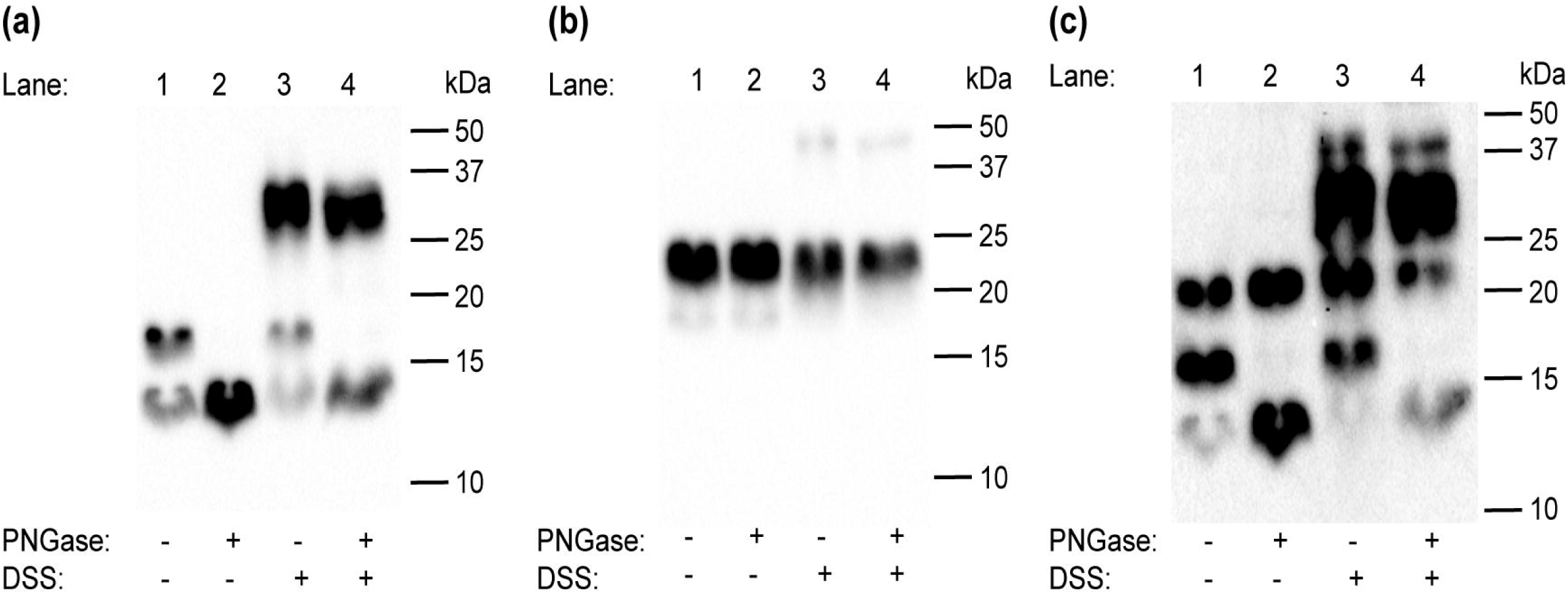
Western blot analyses to determine the effects of glycosylation on homo- and heterodimer formation on the glycoprotein hormone (GPA2/GPB5) subunits in the mosquito, *A. aegypti*. (a) In untreated conditions, western blot analysis of GPA2 subunit alone reveals two bands at 16 and 13 kDa. Whereas following treatment with PNGase, the higher molecular weight band disappears and the 13 kDa band is intensified. A thick, additional band at ∼32 kDa appears when GPA2 protein is cross-linked with DSS, and this band migrates slightly lower to ∼30 kDa when GPA2 protein is treated with both DSS and PNGase. (b) A 24 kDa band is observed in lanes loaded with untreated GPB5 subunit alone. Upon PNGase treatment, the 24 kDa band is not affected; however, upon treatment with DSS, a second faint band appears at 48 kDa that is not affected by deglycosylation. (c) Western blot analyses of co-expressed GPA2 and GPB5 subunits shows three distinct band sizes at 24 kDa, 16 kDa and 13 kDa, corresponding to the GPB5 subunit and two forms of GPA2 subunit protein. Similar to (a), after treatment with PNGase, the higher molecular weight form of GPA2 is eliminated and the 13 kDa band intensifies. When GPA2/GPB5 protein is cross-linked, two additional bands are detected at ∼48 kDa and ∼ 32 kDa; however, following cross-linking and PNGase treatment, the ∼32 kDa band is eliminated leaving only the 30 kDa band along with the unaffected ∼48 kDa band.

### Heterodimerization of mosquito and human GPA2/ GPB5

Using yeast two-hybrid analyses and cross-linking experiments, it was previously shown that human GPA2 (hGPA2) and GPB5 (hGPB5) subunits are capable of heterodimerization ^3^. As a result, to verify whether *A. aegypti* GPA2 and GPB5 subunit proteins are heterodimeric candidates, experiments were first performed alongside hGPA2/hGPB5 subunit proteins, using the latter as an experimental control. Initially, single-promoter expression constructs were designed to incorporate a FLAG-tag and His-tag on the C-terminus of (human and mosquito) GPA2 and GPB5 subunits, respectively (i.e. GPA2-FLAG and GPB5-His). When probed with an anti-His antibody, no bands were detected in lanes containing only GPA2-FLAG (human and mosquito) protein (Fig. 4a, b).

**Fig. 4.**
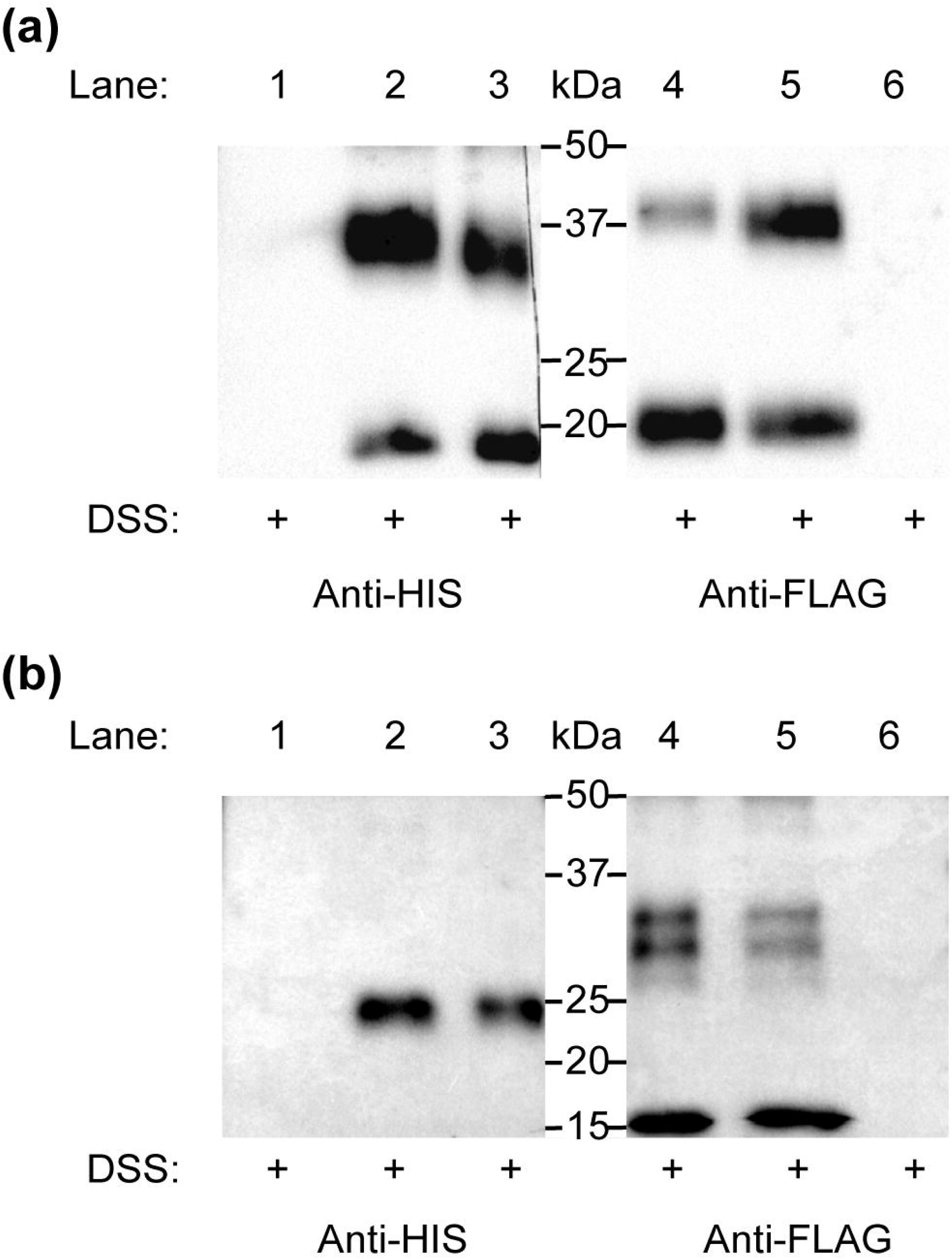
Elucidating heterodimerization of *H. sapiens* (human) (a) and *A. aegypti* (mosquito) (b) GPA2 and GPB5 subunits. Single promoter expression constructs for human and mosquito GPA2-FLAG and GPB5-His were used for transient expression in HEK 293T cells. Protein was harvested and subsequently concentrated, treated with DSS cross-linker, and probed with an anti-His or an anti-FLAG antibody after SDS-PAGE. (a, b) No bands were detected in lanes loaded with cross-linked GPA2-FLAG protein (Lane 1) or cross-linked GPB5-His protein (Lane 6), probed with an anti-His antibody or anti-FLAG antibody, respectively. (a) Bands corresponding to the monomeric form (18 kDa) and homodimer (36 kDa) of cross-linked human GPB5-His (Lane 3) and to the monomeric form (20 kDa) and homodimer (40 kDa) of cross-linked human GPA2-FLAG protein (Lane 4). A combination of the subunits with subsequent cross-linking of separately-produced human GPA2-FLAG and GPB5-His protein (Lane 2, 5) revealed a band size correlating to the human GPA2/ GPB5 heterodimer (38 kDa), detected using an anti-His (Lane 2) or anti-FLAG (Lane 5) antibody. (b) Bands corresponding to the mosquito GPB5 monomer (24 kDa) (Lane 3), mosquito GPA2 glycosylated monomer (16 kDa) and homodimer pairs (30 kDa and 32 kDa) (Lane 4). No detection of bands correlating to mosquito GPA2/GPB5 heterodimer (37-40 kDa) were observed, when probed with either anti-His (Lane 2) or anti-FLAG (Lane 5) antibodies.

Similarly, no bands were detected when an anti-FLAG antibody was used to detect GPB5-His protein (human and mosquito) (Fig. 4a, b). Lanes loaded with cross-linked human GPB5-His protein revealed two bands at 18 kDa (monomer) and 36 kDa (homodimer) (Fig. 4a). Similarly, in lanes loaded with cross-linked human GPA2-FLAG protein, two bands at 20 kDa (monomer) and 40 kDa (homodimer) were detected (Fig. 4a). To determine subunit heterodimerization, GPA2-FLAG protein was combined with GPB5-His protein (i.e. produced separately in different cell batches) and subsequently crosslinked. Results showed an intense 38 kDa band size that correlated to the molecular weight of the hGPA2-FLAG/hGPB5-His heterodimers; detected with both anti-His and anti-FLAG primary antibody solutions (Fig. 4a).

The same experiments were replicated using *A. aegypti* GPA2-FLAG/ GPB5-His protein but using three-fold higher concentration of DSS cross-linker to help improve detection of inter-subunit interactions. Results demonstrated that lanes loaded with DSS-treated GPB5-His protein resulted in a 24 kDa monomer band (Fig. 4b), whereas lanes containing cross-linked GPA2-FLAG detected a 16 kDa (glycosylated monomer), 30 kDa and 32 kDa (homodimers) band size (Fig. 4b). As a result, unlike immunoblots containing hGPA2-FLAG and hGPB5-His protein (Fig. 4a), lanes loaded with cross-linked *A. aegypti* GPA2-FLAG and GPB5-His failed to provide evidence of bands correlating to the predicted molecular weight of an *A. aegypti* GPA2/GPB5 heterodimer, which would be expected at 37-40 kDa (Fig. 4b).

### *A. aegypti* GPA2/GPB5 unable to activate LGR1-mediated Gs and Gi/o signalling pathways

Bioluminescent assays were employed to confirm LGR1 interaction with mosquito GPA2 and/or GPB5 subunits and elucidate downstream signalling pathways upon receptor activation. Since human GPA2/GPB5 has previously been shown to bind and activate human thyrotropin receptor (hTSHR) mediating a stimulatory G protein (Gs) signalling pathway ^3^, our experimental design was first tested using hGPA2/GPB5 and hTSHR as a positive control. Recombinant hGPA2 and hGPB5 subunit proteins were produced in HEK 293T cells with single promoter expression constructs containing the hGPA2 or hGPB5 sequences. Conditioned culture media containing secreted proteins were subsequently concentrated, and crude extracts containing hGPA2, hGPB5 or a combination of hGPA2 and hGPB5 subunits were tested as ligands on HEK 293T cells expressing the hTSHR and a mutant luciferase biosensor, which produces bioluminescence upon interacting with cAMP. Control treatments involved the incubation of hTSHR/ luciferase-expressing cells with concentrated media collected from mCherry-transfected cells (negative control), or 250 nM forskolin (positive control) (Fig. 5a, b). Our findings indicate that incubation with extracts containing a combination of both hGPA2 and hGPB5 were required to stimulate a cAMP-mediated luminescent response from hTSHR-expressing cells, but not when incubated with extracts containing individual subunits (Fig. 5a). We next performed parallel experiments using *A. aegypti* GPA2/GPB5 and LGR1. Given the availability of a dual promoter plasmid (pBud-CE), an additional treatment was performed whereby GPA2/GPB5 subunits were co-expressed within the same cells and conditioned media was concentrated as above. Unlike hGPA2/hGPB5 subunit homologs (Fig. 5a), no combination of mosquito GPA2 and/or GPB5 subunit proteins led to an increase in the luminescent response (Fig. 5b).

**Fig. 5.**
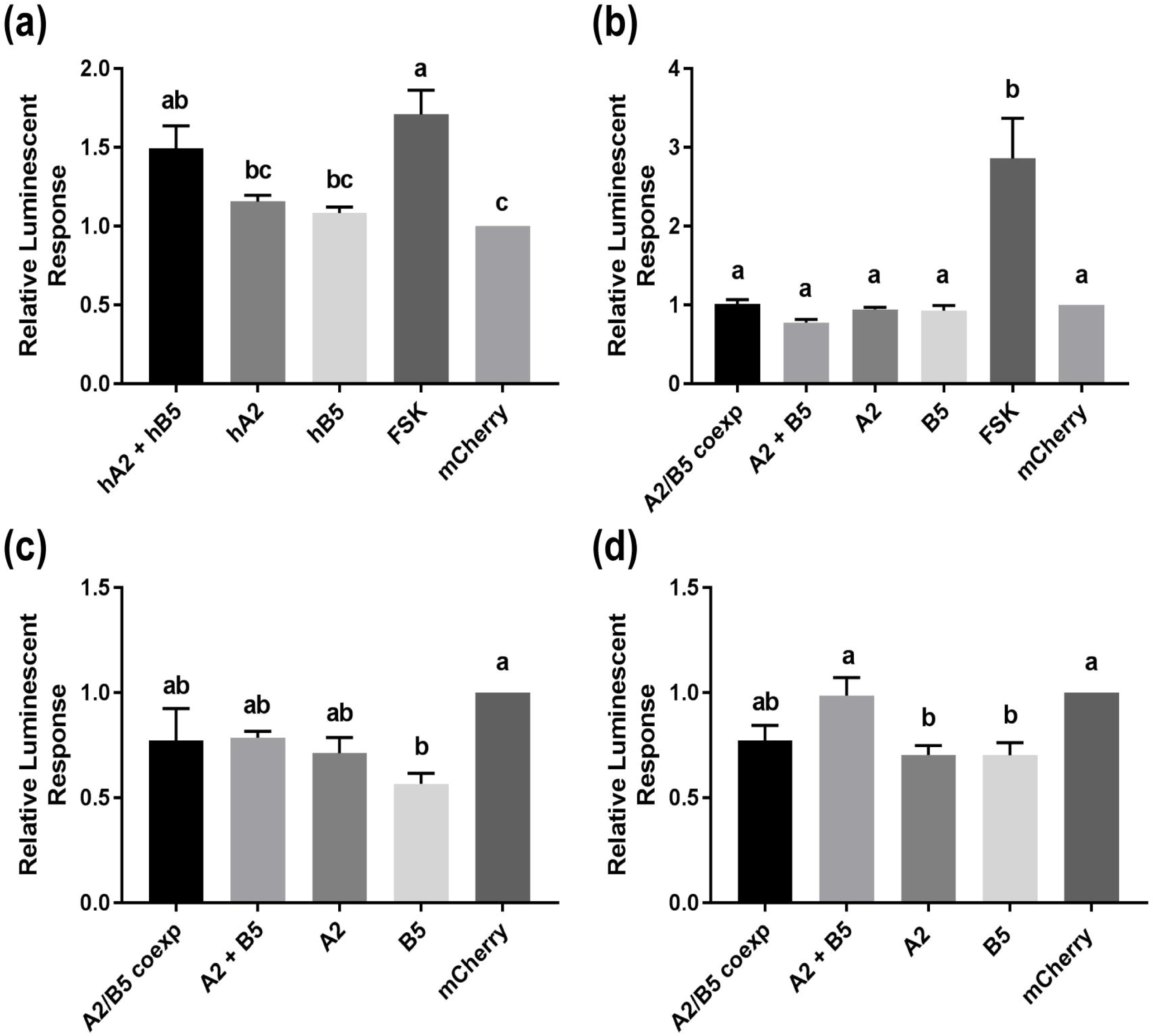
cAMP-mediated bioluminescence assays to determine the effect of GPA2, GPB5 and GPA2/ GPB5 on G-protein signalling of *H. sapiens* (human) TSHR (a), *A. aegypti* (mosquito) LGR1 (b, c), or cells expressing a red fluorescent protein, mCherry (d). Secreted protein fractions for each subunit were prepared separately from HEK 293T cells expressing human GPA2 (hA2), human GPB5 (hB5), mosquito GPA2 (A2), mosquito GPB5 (B5), mCherry, or co-expressing mosquito GPA2 and GPB5 in a dual promoter plasmid (A2/B5 coexp). Secreted protein fractions were then tested separately or combined (A2 + B5) and then incubated with cells co-expressing the cAMP biosensor along with either (a) human TSHR, (b, c) mosquito LGR1 or (d) mCherry, the latter of which was used as a negative control in the functional assay and also served to verify transfection efficiency of HEK 293T cells. Luminescent values were recorded and normalized to incubations with protein secretions collected from the media of mCherry-transfected cells. Forskolin (FSK, 250 nM) was used as a positive control for stimulatory G-protein (Gs) pathway (a, b) and inhibitory G-protein (Gi/o) pathway testing (c, d). (a) Unlike treatments with human GPA2 (hA2) or human GPB5 (hB5) applied singly, a significant increase in cAMP-mediated luminescence was observed when TSHR-expressing cells were incubated with culture media containing both human GPA2 and human GPB5 (hA2 + hB5), relative to incubations with mCherry controls. (b) No differences in luminescence were observed when LGR1-expressing cells were incubated with media containing mosquito GPA2/GPB5 subunits, compared to mCherry controls. (c) The addition of GPA2 and GPB5 on LGR1-expressing cells significantly inhibited FSK-induced luminescent response, compared to treatments with mCherry controls; (d) however, this inhibition was also observed when GPA2 and GPB5 proteins were incubated with HEK 293T cells in the absence of LGR1. Mean ± SEM of three (a, b, d) or six (c) biological replicates. Columns denoted with different letters are significantly different from one another. Multiple comparisons one-way ANOVA test with Tukey’s multiple comparisons (P<0.05).

Using an *in silico* analysis to predict coupling specificity of *A. aegypti* LGR1 and human TSHR to different families of G-proteins ^30^, we determined that *A. aegypti* LGR1 is strongly predicted to couple to inhibitory (Gi) G proteins (Table S3). As a result, to determine whether *A aegypti* GPA2 and/or GPB5 activate a Gi/o signalling pathway, various combinations of GPA2 and GPB5 were tested for their ability to inhibit a forskolin-induced rise in cAMP measured by changes in bioluminescence (Fig. 5c, d). Results revealed that sole treatments of GPA2 and GPB5 proteins alone significantly inhibited a forskolin-induced luminescent response, relative to control treatments with mCherry proteins, when incubated with cells expressing LGR1 (Fig. 5c). However, similar inhibitory effects of GPA2 and GPB5 proteins were also observed with cells not expressing LGR1 (Fig. 5d).

### Characterization of a tethered *A. aegypti* GPA2/GPB5 heterodimer

Activation of hTSHR was only observed when both human GPA2 and GPB5 subunits were coapplied for receptor binding (Fig. 5a) and, unlike the heterodimerization of human GPA2/GPB5 observed in our experiments, mosquito GPA2/GPB5 lacked evidence of heterodimerization (Fig. 4). In light of these observations, it was hypothesized that the activation of *A. aegypti* LGR1 also required subunit heterodimerization. To produce stable GPA2/GPB5 heterodimers using the heterologous expression system, both GPA2 and GPB5 mosquito subunits were expressed as a tethered, single-chain polypeptide by fusing the C-terminus of the GPB5 prepropeptide sequence with the N-terminus of the GPA2 propeptide sequence using a tagged linker sequence composed of twelve amino acids, involving three glycine-serine repeats and six histidine residues.

HEK 293T cells were transfected to transiently express a single promoter plasmid construct containing the tethered GPA2/GPB5 sequence, or the red fluorescent protein (mCherry) as a negative control. At 48-hrs post-transfection, cell lysates along with the conditioned culture media, the latter of which contains secreted proteins, were collected for immunoblot analysis. No bands were detected in lanes containing cell lysate or secreted protein fractions of mCherry transfected cells (Fig. 6a). However, a strong band was detected at 32-40 kDa in the lysates of cells transfected to express tethered GPA2/GPB5 (Fig. 6a). Moreover, two bands were detected at 37 kDa and 40 kDa in lanes loaded with secreted fractions of tethered GPA2/GPB5-transfected cells, that matched the predicted molecular weights of *A. aegypti* GPA2/GPB5 heterodimers corresponding to either non-glycosylated (13 kDa) or glycosylated (16 kDa) GPA2 plus GPB5 (24 kDa) producing bands of 37 kDa or 40 kDa, respectively (Fig. 6a). After secreted protein extracts containing tethered GPA2/GPB5 proteins were treated with PNGase, the higher 40 kDa molecular weight band disappears and the 37 kDa band size intensifies, which confirms the observed molecular weight shift results from removal of N-linked oligosaccharides (Fig. 6b).

**Fig. 6.**
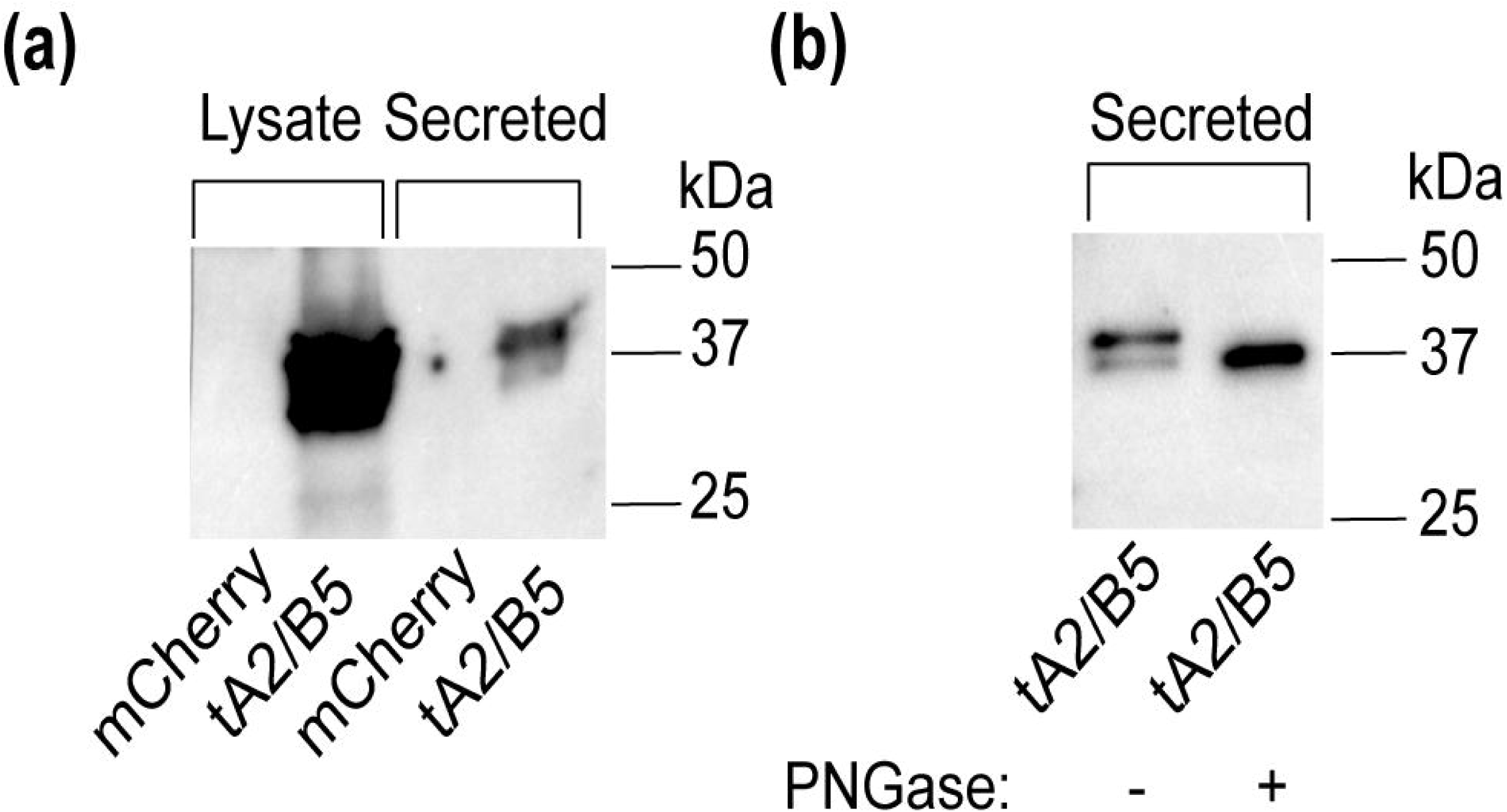
Western blot analysis and verification of *A. aegypti* GPA2/GPB5 tethered protein expressed in HEK 293T cells. (a) Western blot analysis of secreted or cell lysate protein fractions of HEK 293T expressing tethered GPA2/GPB5 (tA2/B5) or red fluorescent protein (mCherry) as a control. Tethered GPA2/GPB5 is represented as a strong 32-40 kDa band in cell lysate fractions, and as two less intense 37 kDa and 40 kDa bands in secreted fractions; however, no bands were detected in lanes loaded with proteins from mCherry transfected cells. (b) Upon treatment of tethered GPA2/GPB5 secreted protein fractions with PNGase, the 40 kDa band is eliminated and the 37 kDa band intensifies, indicating removal of N-linked oligosaccharides.

### *A. aegypti* GPA2/GPB5 heterodimers activate LGR1

The effects of tethered GPA2/GPB5 heterodimers on LGR1 activity was examined. Cell lysates or secreted protein fractions collected from mCherry- or tethered GPA2/GPB5-transfected cells were incubated with HEK 293T cells co-expressing the cAMP luciferase biosensor and either *A. aegypti* LGR1 or mCherry (i.e. not expressing LGR1). Whether tethered GPA2/GPB5 proteins could elevate cAMP or inhibit a forskolin-induced rise in cAMP was assessed and compared to control treatments with proteins harvested from mCherry-transfected cells. Overall, the relative luminescent response was significantly greater in LGR1-transfected cells compared to mCherry-transfected cells (Fig. 7). Unlike treatments with forskolin that significantly increased cAMP-mediated luminescence, neither secreted protein fractions nor cell lysates of tethered GPA2/GPB5-transfected cells elicited an increase in the cAMP-mediated luminescent response relative to controls (Fig. 7a, b). Secreted protein fractions containing tethered GPA2/GPB5 protein had no effect on the forskolin-induced cAMP-mediated luminescence, compared to control treatments with mCherry secreted proteins in LGR1-expressing cells (Fig. 7c). Notably, however, treatments of LGR1-transfected cells, but not mCherry-transfected cells or assay media, with cell lysates containing tethered GPA2/GPB5 protein significantly inhibited the forskolin-induced rise in cAMP relative to treatments with mCherry-transfected cell lysates (Fig. 7d).

**Fig. 7.**
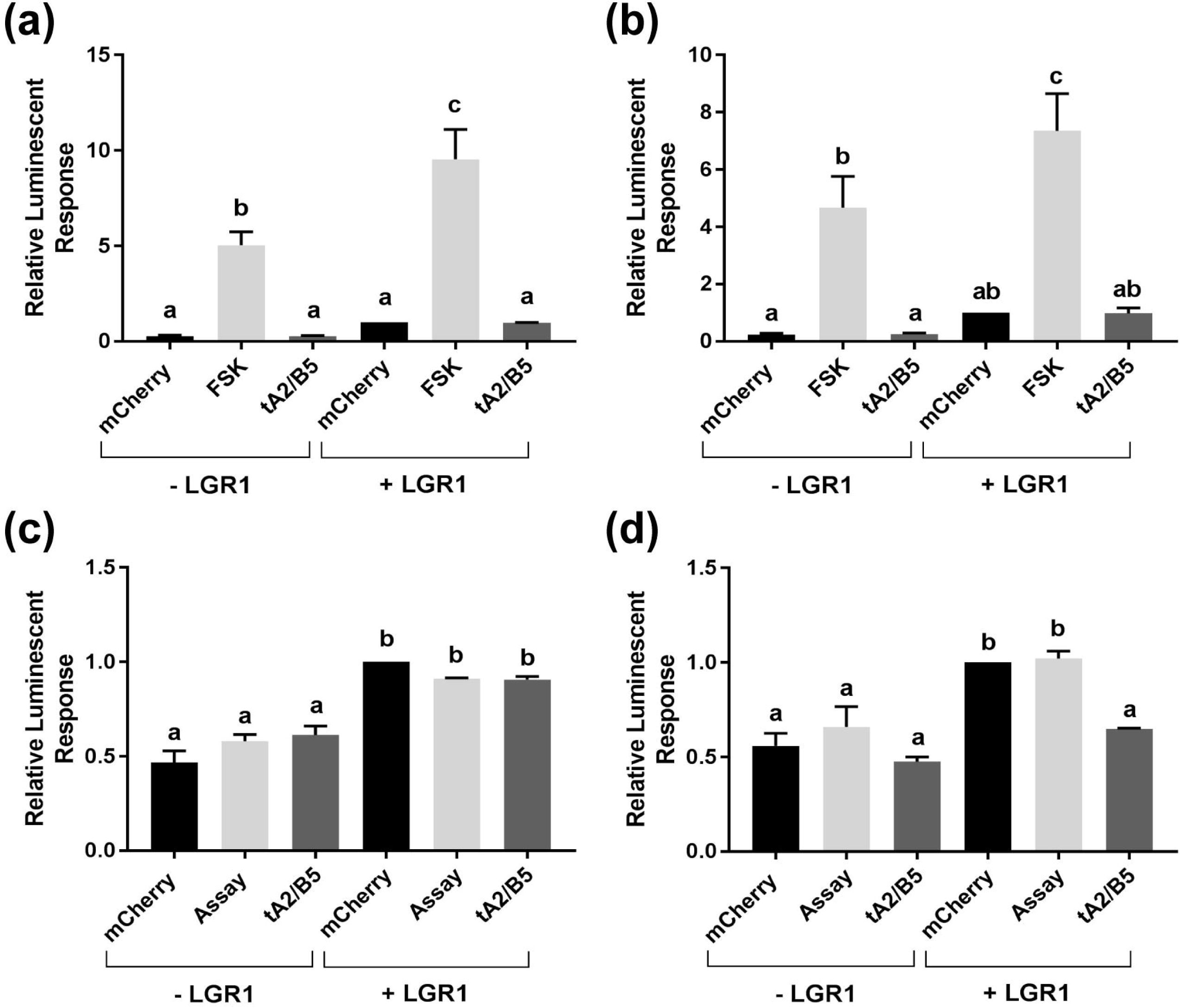
cAMP-mediated bioluminescence assay determining the effects of tethered GPA2/GPB5 on receptor activation and G protein signalling of LGR1. Secreted protein fractions (a, c) and cell lysates (b, d) derived from cells transiently expressing tethered GPA2/GPB5 (tA2/B5) or red fluorescent protein (mCherry) were tested for their ability to stimulate (Gs signalling) (a, b) or inhibit 1 µM forskolin-induced (Gi/o signalling) (c, d) cAMP-mediated luminescence. (a-d) Luminescence response was recorded and normalized to treatments involving mCherry proteins with LGR1-expressing HEK 293T cells. In all treatments, the relative luminescence response was greater in LGR1-expressing cells (+ LGR1) compared to cells not expressing LGR1 (-LGR1). (a, b) Applications of 1 µM forskolin (FSK) to recombinant cells expressing and not expressing LGR1 significantly increased cAMP-mediated luminescence relative to treatments with tA2/B5 or mCherry. However, incubation of LGR1-expressing cells with tA2/B5 secreted (a) or cell lysate (b) proteins, failed to increase cAMP-mediated luminescence above background levels from incubations with mCherry proteins. (c, d) 1 µM forskolin along with either mCherry proteins, tA2/B5 proteins or assay media (Assay) was added to cells in the presence or absence of LGR1 expression. The ability for each ligand treatment to reduce a forskolin-induced increase in cAMP luminescence was examined. (c) The tA2/B5 secreted protein samples incubated with LGR1-expressing cells did not significantly affect the forskolin-induced cAMP luminescence, compared to control treatments with secreted proteins from mCherry expressing cells. (d) Relative to incubations with mCherry cell lysate proteins, cell lysates containing tA2/B5 protein significantly inhibited forskolin-induced elevations of cAMP-mediated luminescence, and this inhibition was specific to LGR1-expressing cells. Mean ± SEM of three biological replicates. Columns denoted with different letters are significantly different from one another. Multiple comparisons two-way ANOVA test with Tukey’s multiple comparisons (P<0.05).

## Discussion

### Co-expression of *A. aegypti* GPA2/GPB5 in neuroendocrine cells of the abdominal ganglia

The central nervous system (CNS) of adult mosquitoes is comprised of a brain and a ventral nerve cord, consisting of three fused thoracic ganglia and six abdominal ganglia. Our findings demonstrate that the GPA2 and GPB5 transcripts are significantly enriched in the abdominal ganglia of adult mosquitoes relative to peripheral tissues and other regions of the CNS, with no sex-specific differences. Although low levels of GPA2 and GPB5 transcripts were detected in the thoracic ganglia and brain using RT-qPCR, fluorescence *in situ* hybridization (FISH) techniques used to localize GPA2 and GPB5 transcripts, along with immunohistochemical detection of GPB5, did not identify specific cells in these regions of the nervous system. Instead, GPA2 and GPB5 transcripts, as well as GPB5 immunoreactivity was identified in 2-3 laterally-localized bilateral pairs of neuroendocrine cells within the first five abdominal ganglia. These findings are consistent with previous findings in the fruit fly *Drosophila melanogaster*, where GPA2 and GPB5 subunit transcripts were localized to four bilateral pairs of neuroendocrine cells in the fused ventral nerve cord, that were distinct from cells expressing other neuropeptides including leucokinin, bursicon, crustacean cardioactive peptide or calcitonin-like diuretic hormone ^16^.

Both FISH and immunohistochemical techniques revealed GPA2 and GPB5 expression within two bilateral pairs of neuroendocrine cells, that were positioned slightly posterior to where the lateral nerves emanate. In some abdominal ganlia preparations, a third bilateral pair of cells immunoreactive for GPB5 protein was detected; however these additional cells were not detected using FISH, which suggests GPB5 transcript may be differentially regulated between different bilateral pairs of cells. Given that the same number of cells were observed to express GPA2 transcript, and these cells localized to similar positions as GPB5 expressing cells, GPA2 and GPB5 were believed to be co-expressed in the same cells. To verify cellular co-expression of GPA2/GPB5, abdominal ganglia were simultaenously treated with both GPA2- and GPB5-targeted anti-sense RNA probes. From this analysis, again only two bilateral pairs of cells were detected and these were more intensly stained compared to preparations treated with either probe alone, which confirms that GPA2 and GPB5 are indeed co-expressed within the same neurosecretory cells of the first five abdominal ganglia in adult mosquitoes. The cellular co-expression of GPA2 and GPB5 proteins implies that, upon a given stimulus, both subunits are regulated in a similar manner and are likely simultaneously released following the appropriate stimulus. Importantly, since co-expression and heterodimerization of the classic vertebrate glycoprotein hormone subunits takes place within the same cells ^21, 22^, these findings indicate that the mosquito GPA2/GPB5 subunits may be produced and released as heterodimers *in vivo*.

### Heterodimerization and Homodimerization of GPA2/GPB5

To study the interactions of *A. aegypti* GPA2 and GPB5 subunits *in vitro*, hexa-histidine tagged proteins secreted into the culture media of transfected HEK 293T cells were collected and analysed under denaturing conditions after cross-linker treatments, which had been utilized previously to show GPA2/GPB5 heterodimerization in other organisms ^3, 6, 29^. Cross-linked protein samples were then deglycosylated to identify whether the removal of N-linked sugars affected dimerization. Treatment of mosquito GPA2 and GPB5 subunits individually with cross-linker resulted in the detection of bands with sizes corresponding to homodimers of GPA2 (∼32 kDa) and GPB5 (48 kDa). GPA2 homodimer bands migrated lower to ∼30 kDa following deglycosylation treatment with PNGase. However, experiments performed with cross-linked GPA2/GPB5 protein were not able to confirm heterodimeriation since the detected band sizes could also reflect GPA2 and GPB5 homodimeric interactions. Previous cross-linking studies that demonstrated GPA2/GPB5 heterodimerization in human ^3, 6^ and fruit fly ^29^ did not provide evidence on the interactions of each subunit alone to determine if homodimerization is possible. In these earlier studies, molecular weight band sizes that were identified as heterodimers could have been the result of homodimeric interactions ^3, 6, 29^. As a result, the findings herein with mosquito GPA2/GPB5 indicate additional experiments are required to confirm GPA2/GPB5 heterodimerization in these organisms.

To clarify whether *A. aegypti* (mosquito) GPA2/GPB5 heterodimerize, each subunit was differentially tagged (GPA2-FLAG and GPB5-His), and immunoblots containing various combinations of cross-linked subunits were probed with either anti-FLAG or anti-His antibody. As a positive control, experiments were first performed using *H. sapiens* (human) GPA2/GPB5 subunit proteins. Similar to mosquito GPA2 and GPB5 subunits, results showed human GPA2 and GPB5 subunits are capable of homodimerization. To study heterodimeric interactions, GPA2 and GPB5 subunit proteins were expressed separately in HEK 293T cells. Upon combining and treating protein samples with DSS, a molecular weight band size at 38 kDa, corresponding to the molecular weight of GPA2/GPB5 heterodimers (human GPA2-FLAG/ GPB5-His), was detected and migrated differently than bands corresponding to GPA2 (40 kDa) and GPB5 (36 kDa) homodimers. Taken together, our results confirm human GPA2 and GPB5 is indeed capable of heterodimerization *in vitro*. Further, an induction of cAMP was observed when both subunits were present for TSHR functional activation, whereas treatments with individual subunits failed to significantly increase cAMP-mediated luminescence. As a result, human GPA2/GPB5 is capable of heterodimerization, and since a combination of both subunits were required to signal a TSHR-mediated elevation in cAMP, these heterodimers are required to functionally activate its cognate glycoprotein hormone receptor (TSHR).

The classic glycoprotein hormones, which include FSH, LH, TSH and CG, are formed by a common alpha subunit non-covalently linked to a hormone-specific beta subunit. Each subunit is co-expressed and assembled as a heterodimer in the endoplasmic reticulum within cells before being secreted as heterodimer into circulation ^21^. Heterodimerization is required for secretion and receptor binding activity of each hormone ^1, 31, 32^. Our results confirm human GPA2/GPB5 heterodimerizes, as detected following chemical cross-linking treatments and, even without cross-linking, both subunits are required to activate TSHR. In our experimental approach, human GPA2 and GPB5 subunits were not co-expressed within the same cells using a dual promoter expression construct. Rather, subunits were separately expressed by different batches of cells and mixed post harvesting of conditioned media to demonstrate heterodimerization and TSHR activation. Thus, unlike FSH, TSH, CG, and LH ^1^, our results provide novel information that demonstrate co-expression is not necessarily required for heterodimerization of human GPA2/GPB5 *in vitro*.

Analogous experiments using the same concentration of DSS cross-linker (data not shown) as well as three-fold higher concentrations were performed with mosquito GPA2-FLAG and GPB5-His subunit proteins. Irrespective of the DSS concentration used, no bands corresponding to the expected molecular weights of GPA2/GPB5 heterodimers (37-40 kDa) were observed, but rather, only GPA2 and GPB5 homodimers were detected (Fig. 4b). The inability of mosquito GPA2 and GPB5 subunits to successfully heterodimerize could result from improper protein folding of insect-derived secretory proteins in HEK 293T cells used for heterologous expression. As a result, future experiments should test the heterologous expression and heterodimerization of *A. aegypti* GPA2/GPB5 in insect cell lines that could provide a more appropriate environment for tertiary and quaternary protein structure formation.

Mosquito GPA2/GPB5 (each subunit alone, mixed from different cell batches or co-expressed in the same cells using a dual promoter vector) was incapable of stimulating a cAMP-mediated luminescent response in HEK 293T cells expressing LGR1. Since we identified *A. aegypti* LGR1 is predicted to couple a Gi/o signalling pathway (Table S3), we next examined if mosquito GPA2/GPB5 could inhibit a forskolin-induced cAMP response. Sole treatments of GPA2 and GPB5 subunit proteins inhibited a forskolin-induced rise in cAMP, however these inhibitory actions were not owed to G protein signalling events related to *A. aegypti* LGR1, since inhibition was observed in control cell lines in the absence of LGR1. These results suggest mosquito GPA2 and GPB5 subunit proteins may non-specifically interact with other endogenously expressed proteins, like the orphan glycoprotein hormone receptors LGR4 and LGR5 that are highly expressed in HEK 293T cells ^33^.

### *A. aegypti* GPA2/ GPB5 heterodimers activate LGR1 and initiate a switch from Gs to Gi coupling

Activation of the human thyrotropin receptor was only observed when both human GPA2 and GPB5 subunits were present which, unlike data obtained involving mosquito subunits, demonstrated human GPA2/GPB5 subunit heterodimerization *in vitro* using the mammalian heterologous system. Nonetheless, given the observed co-localization of the subunits within bilateral pairs of cells in the first five abdominal ganglia in *A. aegypti*, which is comparable to cellular colocalization shown earlier in *D. melanogaster* ^16^, we proposed that *A. aegypti* GPA2/GPB5 heterodimers would be required to functionally activate LGR1 *in vitro*. To confirm this possibility, a tethered construct was designed linking the C-terminus of GPB5 to the N-terminus of GPA2 using a histidine tagged glycine/serine-rich linker sequence. Natural and synthetic linkers function as spacers that connect multidomain proteins, and are commonly used to study unstable or weak protein-protein interactions ^34^. The incorporation of a linker sequence between glycoprotein hormone subunits has been performed previously, and does not affect the assembly, secretion or bioactivity of human FSH ^35^, TSH ^36^ and CG ^37^. The conversion of two independent glycoprotein hormone subunits into a single polypeptide chain using a glycine-serine repeat linker sequence has also been performed recently with lamprey GPA2/GPB5 ^38^, which was shown to induce a cAMP response. Interestingly, similar proteins involving TSH alpha fused to TSH beta with carboxyl-terminal peptide (CTP) as a linker promoted a three-fold higher induction of cAMP compared to wild-type TSH, likely because the addition of a CTP linker increases protein stability and flexibility ^39^.

Thus, tethered *A. aegypti* GPA2/GPB5 was expressed in HEK 293T cells and cell lysates and secreted protein fractions were collected for expression studies. Immunoblot studies revealed the detection of a strong 32-40 kDa band in lanes loaded with cell lysate proteins, and two less intense distinct bands at 40 kDa and 37 kDa in secreted protein fractions of GPA2/GPB5-transfected cells. Molecular weight band sizes at 40 kDa and 37 kDa in secreted protein fractions matched the expected band sizes of tethered GPA2/GPB5 heterodimers, corresponding to nonglycosylated 13 kDa or glycosylated 16 kDa GPA2 plus 24 kDa GPB5. Moreover, treatments with PNGase verified the tethered GPA2/GPB5 proteins are glycosylated, as observed for GPA2 expressed independantly in earlier experiments herein and in previous studies ^14^. Highly intense bands in cell lysates indicate the retention of immature and mature GPA2/GPB5 heterodimers. For this reason, cell lysates and secreted protein fractions of tethered GPA2/GPB5-expressing cells were individually tested for their ability to activate *A. aegypti* LGR1.

In humans, GPA2/GPB5-TSHR signalling stimulates adenylyl cyclase activity to increase intracellular cAMP via interaction with a Gs protein ^3, 6, 11^, and these results were confirmed in our studies. Comparatively, GPA2/GPB5 signalling was also shown to increase levels of cAMP upon binding LGR1 in *D. melanogaster* ^29^. Interestingly, our experiments demonstrate low level constitutive activity of adenylyl cyclase in LGR1 expressing cells since cAMP luminescent response was moderately greater in LGR1-transfected cells compared to cells not expressing LGR1. Constitutive activity of glycoprotein hormone receptors is well known and has been demonstrated to be stronger for the thyrotropin receptor than for the LH/CG receptor ^40^. Surprisingly, our experiments indicate that incubations of LGR1-expressing cells with GPA2/GPB5 tethered heterodimers triggers a switch from low level constitutive Gs coupling to Gi coupling for *A. aegypti* LGR1, given that GPA2/GPB5 heterodimers signficantly inhibited the forskolin-induced increase in cAMP in LGR1-transfected cells but not in cells lacking LGR1 expression. This finding, while highly interesting, is not entirely unusual since promiscuous G protein coupling has been reported for glycoprotein hormone receptors like the TSH receptor (Gs and Gq) and LH/CG (Gi and Gs) ^41–43^.

### Regulation by GPA2/GPB5 heterodimers

To help stabilize heterodimerization, the beta subunit sequences of the classic glycoprotein hormones (FSH, LH, TSH and CG) contain two additional cysteine residues that form an additional disulfide bridge which wraps around and “buckles” the alpha subunit ^26^. Though heterodimerization can occur with mutated forms of this “seatbelt” structure, there is a dramatic decrease in heterodimer stability ^21, 22^. GPB5 in vertebrates and invertebrates lack the seatbelt structure required to stabilize heterodimerization ^26^. Thus, the hypothesis that GPA2/ GPB5 functions as a heterodimer in a physiological situation (i.e. without chemical cross-linking) is challenged. The dissociation constant (Kd) associated with heterodimerization of the classic glycoproteins hormone subunits and GPA2/GPB5 is 10^-7^ M to 10^-6^ M, which indicates heterodimeric interactions are favoured at these concentrations ^44, 45^. However, since the classic beta subunits contain an additional disulfide bridge that strengthens its association with the common alpha subunit, heterodimeric interactions are stabilized in circulation at physiological concentrations as low as 10^-11^ M to 10^-9^ M ^22^. Without this seatbelt structure, GPA2/GPB5 heterodimeric interactions are posssible only at micromolar concentrations, which are not typically observed in circulation ^22, 26^. Together, the limited evidence so far using heterolougous expression challenges the possibility of endocrine regulation by GPA2/GPB5 heterodimers. Nonetheless, it was argued in *D. melanogaster* that the large neurosecretory cells co-expressing the glycoprotein hormone subunits, along with their corresponding axonal projections that localized distinctly from organs that express the GPA2/GPB5 receptor (LGR1), did support that this system is indeed endocrine in nature ^16^. Alternatively, the subunits could function independently or regulate physiology as a heterodimer in a paracrine/ autocrine fashion. In rats, GPA2/GPB5 is expressed in oocytes and may act as a paracrine regulator of TSHR-expressed granulosa cells in the ovary to regulate reproductive processes ^6^. Another possibility to consider is that additional endogenous co-factors may be involved, but remain unidentified, which help to strengthen interaction between the GPA2 and GPB5 subunits, since the tethered mosquito GPA2/GPB5 was indeed capable of activating LGR1 *in vitro* inducing a Gi signalling cascade.

### GPA2 and GPB5 homodimerization

Our results establish that human and mosquito GPA2 and GPB5 subunits can weakly and strongly, respectively, homodimerize. However, whether these homodimers, pertain to a physiological function *in vivo* is unknown. Treatments of either mosquito or human GPA2 and GPB5 subunits alone did not stimulate specific downstream signalling in LGR1 or TSHR-expressing cells, indicating that only GPA2/GPB5 heterodimers can activate their cognate glycoprotein hormone receptors. However, it is possible that GPA2 and GPB5 homodimers may target other unidentified receptors. In insects, the moulting hormone bursicon is a heterodimer of two subunits called burs and pburs. Burs/pburs heterodimers act via a glycoprotein hormone receptor (i.e. LGR2) to regulate processes such as tanning and sclerotization of the insect cuticle as well as wing inflation after adult emergence ^46^. Recently, it was demonstrated that bursicon subunits can homodimerize (i.e. burs/burs and pburs/pburs) and these homodimers mediate actions independently of LGR2 to regulate immune responses in *A. aegypti* and *D. melanogaster* ^47, 48^.

In addition to the human/ mosquito GPA2/GPB5 homodimers observed in our studies, human GPA2 was also shown to interact with the beta subunits of CG and FSH ^3^. Lastly, the expression patterns of GPA2 and GPB5 in a number of organisms do not always strictly co-localize, since GPA2 expression exhibits a much wider distribution and is expressed more abundantly than GPB5 in a number of vertebrate and invertebrate organisms ^7, 12, 23, 24, 49, 50^. Taken together, this raises the possibility that GPA2 and GPB5 subunits may interact with other unknown proteins that could activate different receptors or signalling pathways and elicit distinct functions.

### Concluding remarks

Although much is known about the classic vertebrate glycoprotein hormones including LH, FSH, TSH and CG along with their associated receptors, little progress has been made thus far towards better understanding the function of GPA2/GPB5, signalling and subunit interactions, particularly for the invertebrate organisms. To our knowledge, this is the first study to demonstrate *A. aegypti* and *H. sapiens* GPA2 and GPB5 subunit homodimerization *in vitro*. Our results also confirm that heterodimerization of *A. aegypti* and *H. sapiens* GPA2/GPB5 are required for the activation of their cognate receptors LGR1 and TSHR, respectively. Unlike previous reports showing GPA2/GPB5-induced LGR1 activation elevates intracellular cAMP by coupling a Gs pathway, the current findings provide novel information supporting that *A. aegypti* LGR1 couples to a Gi protein to inhibit cAMP levels following application of heterodimeric GPA2/GPB5. Further, our results revealed that mosquito LGR1 is constitutively active when overexpressed in the absence of its ligand, GPA2/GPB5, inducing a Gs signalling pathway that raises levels of cAMP levels, which is consistent with previous observations with overexpression of fruit fly LGR1 ^29, 51^ as well as mammals including dog and human TSH receptor ^52, 53^.

In the mosquito nervous system, GPA2 and GPB5 subunits are co-expressed within the the same neurosecretory cells of the first five abdominal ganlgia where their coordinated release and regulation are likely. As a result, whether GPA2/GPB5 are secreted as heterodimers, like the classic glycoprotein hormones, and/or as homodimers remains to be determined *in vivo*; however, these results confirm GPA2 and GPB5 homodimers do not activate LGR1 and TSHR. Whether these homodimers are functional *in vivo* and what physiological role they play (if any) is a research direction that should be addressed in future studies. All in all, this investigation has provided novel information for an invertebrate GPA2/GPB5 and LGR1 signalling system and contributes towards advancing our understanding and the functional elucidation of this ancient glycoprotein hormone signalling system common to nearly all bilaterian organisms.

## Acknowledgements

Research in this study was supported by a Natural Sciences and Engineering Research Council of Canada (NSERC) Discovery Grant and Ontario Ministry of Research & Innovation Early Researcher Award to JPP.

## Materials and methods

### Animals

Adult *Aedes aegypti* (Liverpool) were derived from an established laboratory-reared colony raised under conditions described previously ^18^.

### GPA2/ GPB5 transcript analysis by RT-qPCR

Total RNA was isolated and purified from select *A. aegypti* tissues and organs, reverse transcribed into cDNA. GPA2 and GPB5 transcript abundance was quantified using a StepOnePlus^TM^ Real-Time PCR System (Applied Biosystems) as described previously ^55^ (see SI Materials and Methods for details). GPA2 and GPB5 transcript abundance was normalized to the expression of two reference genes following the ΔΔCt method as previously described ^14^. Experiments were repeated using a total of three technical replicates per sample and three biological replicates for each tissue/ organ.

### Fluorescence in situ hybridization

Using gene-specific primers (Table S1), *A. aegypti* GPA2 and GPB5 sequences were amplified from previously prepared constructs ^14^ that contained the GPA2 and GPB5 complete open reading frames (ORFs). Sense and antisense probes were then generated following a similar protocol as recently reported ^19^. Briefly, cDNA fragments were ligated to pGEM T Easy vector (Promega, Madison, WI, USA) and used to transform NEB 5-α competent *Escherichia coli* cells (New England Biolabs, Whitby, ON, Canada). After screening plasmid constructs for directionality using T7 promoter oligonucleotide and gene-specific primers (Table S1), template sense or anti-sense cDNA strands for GPA2 and GPB5 probe synthesis were created by PCR amplification and verified by Sanger sequencing for base accuracy (The Centre for Applied Genomics, Sick Kids Hospital, Toronto, ON, Canada). Digoxigenin (DIG)-labelled anti-sense and sense RNA probes corresponding to GPA2 and GPB5 subunits were synthesized using the HiScribe T7 High Yield RNA Synthesis kit (New England Biolabs, Whitby, ON, Canada). Fluorescence *in situ* hybridization (FISH) was then used to detect GPA2 and/or GPB5 transcript in the mosquito central nervous system using 4 ng µl^-1^ (GPB5) and/or 6 ng µl^-1^ (GPA2) RNA sense/ antisense probes, following a previously established protocol ^54^. Preparations were analyzed with a Lumen Dynamics X-CiteTM 120Q Nikon fluorescence microscope (Nikon, Mississauga, ON, Canada), or a Yokogowa CSU-XI Zeiss Cell Observer Spinning Disk confocal microscope, and images were processed using Zeiss Zen and ImageJ software. All microscope settings were kept identical when acquiring images of control and experimental preparations.

### Wholemount immunohistochemistry

GPB5 immunoreactivity in the abdominal ganglia of adult mosquitoes was examined in newly emerged and four-day old *A. aegypti* that were lightly CO_2_ anesthetized, and dissected in PBS at RT. Tissues were fixed, permeabilized and incubated in a custom rabbit polyclonal GPB5 antibody (1 μg ml^−1^) designed against an antigen sequence (CDSNEISDWRFP) positioned at residues 69-80 ^14^ for 48 h at 4°C rocking. Control treatments involved pre-incubated *A. aegypti* GPB5 primary antibody solution containing 100:1 peptide antigen:antibody (mol:mol). After several washes, tissues were incubated overnight at 4°C in Alexa Fluor 488-conjugated goat anti-rabbit Ab (1:200) secondary antibody (Life Technologies) in PBS containing 10% normal sheep serum. The next day, samples were washed and then mounted onto coverslips using mounting media containing Diamidino-2-phenylindole dihydrochloride (DAPI), and analyzed using a Lumen Dynamics X-CiteTM 120Q Nikon fluorescence microscope (Nikon, Mississauga, ON, Canada), or optically sectioned using a Yokogowa CSU-XI Zeiss Cell Observer Spinning Disk confocal microscope. All images were processed using Zeiss Zen and ImageJ software. Further details concerning the wholemount immunohistochemical protocol were reported in an earlier study ^18^.

### Plasmid expression constructs

Plasmid expression constructs were designed to study *A. aegypti* and *H. sapiens* GPA2/GPB5 subunit dimerization patterns and receptor signalling. Using previously available hexa-histidine-tagged *A. aegypti* GPA2-His ^14^, as well as FLAG (DYKDDDDK)-tagged (-FLAG) *H. sapiens* GPB5-FLAG (Genscript, Clone OHu31847D) plasmid vectors as template, the full ORF of each (*A. aegypti* and *H. sapiens*) GPA2 and GPB5 subunit coding sequence, including a consensus Kozak translation initiation sequence, was amplified and a hexa-histidine or FLAG tag sequence was incorporated on the carboxyl-terminus of subunits to produce the following fusion proteins; *A. aegypti* GPA2-FLAG, *H. sapiens* GPB5-His (Table S2). A pcDNA3.1^+^ mammalian expression construct containing mCherry, which was a gift from Scott Gradia (Addgene plasmid # 30125), was utilized to verify cell transfection efficiency. Experiments also utilized previously prepared pcDNA3.1^+^ constructs with *A. aegypti* GPB5-His and *A. aegypti* LGR1 coding sequences and dual promoter vector pBudCE4.1 containing both *A. aegypti* GPA2-His and GPB5-His ^55^. Additionally, pcDNA 3.1^+^ mammalian expression vector construct containing FLAG tagged *H. sapiens* thyrotropin receptor (TSHR-FLAG) (Genscript USA Inc., Clone OHu18318D), *H. sapiens* GPA2-FLAG (Genscript USA Inc., Clone OHu31847D), *H. sapiens* GPB5-FLAG (Genscript USA Inc., Clone OHu55827D) and pGlosensor^TM^ -22F cyclic adenosine monophosphate (cAMP) biosensor plasmid (Promega Corp., Madison, WI), which were used for receptor activation and intracellular signalling assays.

### Generation of tethered *A. aegypti* GPA2/GPB5 construct

The ORFs of *A. aegypti* GPA2 and GPB5 sequences were tethered together in order to promote heterodimer interactions for testing in receptor activity assays with mammalian cell lines. A hexa-histidine tagged artificial linker sequence involving three glycine-serine repeats was used to fuse the amino-terminus of *A. aegypti* GPA2 propeptide sequence to the carboxyl-terminus of *A. aegypti* GPB5 prepropeptide sequence, using multiple PCR amplifications with several primer sets (Table S2) as performed previously using lamprey GPA2 and GPB5 sequences ^38^ (see SI Materials and Methods for details).

### Transient transfection of HEK 293T cells

Human embryonic kidney (HEK 293T) cells were grown in complete growth media (Dulbecco’s modified eagles medium: nutrient F12 (DMEM) media, 10% heat inactivated fetal bovine serum (Wisent, St. Bruno, QC) and 1X antimycotic-antibiotic) and maintained in a water-jacketed incubator at 37°C, 5% CO_2_. When cells reached ∼80-90% confluency, they were transfected with plasmid expression constructs in 6-well tissue culture plates (Thermo Fisher Scientific, Burlington, ON) using Lipofectamine 3000 transfection reagent (Life Technologies, Carlsbad, CA) with 3:1 (µL:µg) transfection reagent to DNA ratio. Before transfection, culture media was replaced with either serum-free medium (DMEM and 1X antimycotic-antibiotic) for experiments that collected secreted proteins, or fresh complete growth medium for experiments that dually-transfected cells with pGlosensor^TM^ -22F cAMP biosensor plasmid and either *H. sapiens* TSHR, *A. aegypti* LGR1, or mCherry.

### Preparation of protein samples

At 48 h post-transfection, serum-free culture media containing secreted proteins were collected and concentrated in 0.5 mL 3-kDa molecular weight cut-off centrifugal filters (VWR North America). In some experiments, cells were dislodged with PBS containing 5 mM ethylenediaminetetraacetic acid (EDTA; Life Technologies) (PBS-EDTA) pH 8.0, pelleted at 400x*g* for 5 min, resuspended in PBS and transferred to 1.5 mL centrifuge tubes for a subsequent centrifugation. Cell lysates were then prepared by resuspending and sonicating cells for 5 s in cell lysis buffer containing 37.5 mM Tris, pH = 7.5, 1.5 mM EDTA, pH 8.0, 3% sodium dodecyl sulfate, 1.5% protease inhibitor cocktail (v/v), and 1.5 mM dithiothreitol (DTT). For receptor activity assays, in order to prevent carry-over of lysis buffer, cell lysates were concentrated in 3-kDa molecular weight cut-off centrifugal filters and re-constituted back to initial volumes with serum-free media for a total of three repetitions. Proteins were then used for cross-linking analysis, deglycosylation and immunoblotting or used as ligands for functional receptor activation using the cAMP signalling biosensor assays.

Dissuccinimidyl suberate (DSS), a chemical cross-linker that is primarily reactive towards amino groups providing stabilization of weak or transient protein intermolecular interactions, was employed to study GPA2 and GPB5 protein-protein interactions. *A. aegypti* and *H. sapiens* GPA2-FLAG and GPB5-His proteins were separately, or combined together (GPA2-FLAG/GPB5-His) and then treated with DSS (Sigma Aldrich, Oakville, ON). In some experiments, *A. aegypti* GPA2-His/GPB5-His were co-expressed using a dual promoter plasmid and as such, media containing both His-tagged subunits were directly treated with DSS. To treat secreted concentrates of culture media containing *H.* sapiens GPA2 and GPB5 proteins, 0.68 mM DSS was used whereas both 0.68 mM (data not shown) and 2.04 mM DSS was used to test the dimerization characteristics of *A. aegypti* GPA2-His/GPB5-FLAG proteins. Cross-linking was performed for 30 min at RT and reactions were quenched with 50 mM Tris, pH 7.4 for 10 min under constant mixing. To remove N-linked sugars, protein samples were treated with peptide-N-Glycosidase F (PNGase) (New England Biolabs, Whitby, ON) following manufacturers guidelines, with the only modification being that protein samples were not heated to 100°C before enzymatic deglycosylation. Experiments aimed to determine the effects of protein glycosylation on cross-linking ability first treated *A. aegypti* GPA2-His/ GPB5-His subunits with PNGase and subsequently cross-linked samples with DSS after deglycosylation.

### Western blot analysis

Samples were prepared in 2x Laemmli buffer (Sigma Aldrich, Oakville, ON) containing 4% SDS, 20% glycerol, 10% 2-mercaptoethanol, 0.004% bromophenol blue and 0.125 M Tris HCl, pH ∼6.8, and resolved on 10% or 15% SDS-polyacrylamide gels under reducing conditions at 120 V for 90-110 min. Using a wet transfer system, proteins were then transferred to polyvinylidene difluoride (PVDF) membranes at 100 V for 75 min. For GPA2-FLAG/GPB5-His heterodimerization experiments, samples were run in duplicate on the same gel and following transfer, membranes were cut in half for separate primary antibody incubations. Membrane blots containing protein samples were blocked for 1 h in PBS containing 0.1% Tween-20 (Bioshop, Burlington, ON, Canada) and 5% skim milk powder (PBSTB) rocking at RT. After blocking, membranes were incubated overnight at 4°C on a rocking platform in PBSTB containing mouse monoclonal anti-His (1:500 dilution) or mouse monoclonal anti-FLAG (1:500) primary antibody solutions. The next day, membranes were washed three times with PBS containing 0.1% Tween-20 (PBST) for 15 min each wash and then were incubated in PBSTB containing anti-mouse HRP conjugated secondary antibody (1:2000 – 1:3000 dilution) for 1 h rocking at RT before washing again with PBST (3 x 15 min washes). Finally, blots were incubated with Clarity Western ECL substrate and images were developed using a ChemiDoc MP Imaging System (Bio-Rad Laboratories, Mississauga, ON) and molecular weight measurements and analysis were performed using Image Lab 5.0 software (Bio-Rad Laboratories, Mississauga, ON).

### Receptor Functional Activation Bioluminescence Assays

HEK 293T cells were co-transfected to express (i) *H. sapiens* TSHR, *A. aegypti* LGR1, or mCherry along with (ii) pGlosensor^TM^ -22F cAMP biosensor plasmid (Promega Corp., Madison, WI), which encodes a modified form of firefly luciferase with a fused cAMP binding moiety providing a biosensor for the direct detection of cAMP signalling in live cells. At 48 h post transfection, recombinant cells were dislodged with PBS-EDTA, pelleted at 400x*g* for 5 min, resuspended in assay media (DMEM:F12 media with 10% fetal bovine serum (v/v)) containing cAMP GloSensor reagent (2% v/v), and incubated for 3 h rocking at RT shielded from light. White 96-well luminescence plates (Greiner Bio-One, Germany) were loaded under low light with previously prepared secreted or cell lysate protein concentrates (described above), forskolin or assay media alone, and incubated at 37°C for 30 min prior to performing receptor activity assays. For stimulatory G-protein (Gs) pathway detection, recombinant cells expressing either *H. sapiens* TSHR, *A. aegypti* LGR1, or mCherry along with the cAMP biosensor were pre-treated with 0.25 mM 3-isobutyl-1-methylxanthine (IBMX) for 30 min at RT with rocking and shielded from light. Using an automatic injector unit (BioTek Instruments Inc., Winooski VT), recombinant cells were then loaded into wells containing various treatments, including 250 nM forskolin as a positive control or concentrates of protein fractions from mCherry-transfected cells as a negative control. For inhibitory G-protein (Gi/o) pathway testing, *A. aegypti* GPA2 and/or GPB5 was tested for the ability to inhibit a forskolin-induced cAMP-mediated bioluminescent response. Using an automatic injector unit (BioTek Instruments Inc., Winooski VT), cells expressing LGR1 or mCherry (LGR1 activation and signalling negative control) were loaded into wells that contained various ligand treatments. Subsequently, 250 nM or 1 µM forskolin was added to wells immediately after the addition of recombinant cells, using a second automatic injector unit. For bioluminescent assays with tethered GPA2/GPB5 proteins, after the 3 h incubation, recombinant cells expressing or not expressing LGR1 were equally divided to test Gs and Gi signalling pathways simultaneously using the same batch of cells per biological replicate. In all assays, luminescence was measured every 2 min for 20 min at 37°C using a Synergy 2 Multimode Microplate Reader (BioTek, Winooski, VT) and averaged over 4-8 technical replicates for each treatment. To calculate the relative luminescent response, data was normalized to luminescent values recorded from negative control ligand treatments involving protein secretions derived from mCherry-transfected cells. Assays were performed repeatedly and involved 3-6 independent biological replicates.

## Notes

#### Summary of Updates

Updated manuscript format and referencing style.

