## Supplementary Information for "Expression Profiling, Downstream Signaling and Subunit Interactions of GPA2/GPB5 in the Adult Mosquito *Aedes aegypti*"

**This PDF file includes:**

SI Materials and Methods

Tables S1, S2 and S3

References for SI-S3 citations

**SI Materials and Methods**

**GPA2/ GPB5 transcript analysis by RT-qPCR**

Male and female adult mosquitoes four-days post-eclosion were lightly CO_2_ anesthetized to allow wing and leg removal and dissected in nuclease-free Dulbecco’s phosphate-buffered saline (PBS; pH 7.3) (Wisent, St. Bruno, QC, Canada) at RT. Dissected tissues and organs included the brain, thoracic ganglia, abdominal ganglia, alimentary canal (including the midgut, hindgut and Malpighian tubules), reproductive organs (including accessory reproductive tissues and ovaries/testes) and carcass (containing peripheral fat body and musculature) of each sex. After dissection, tissues were immediately placed into RNA lysis buffer (Bio Basic, Markham, ON, Canada) and stored overnight at -20°C. Upon thawing samples at RT, total RNA was purified using the EZ-10 RNA Mini-Preps Kit (Bio Basic, Markham, ON, Canada) that included an on-column DNase I treatment to remove potential contaminating genomic DNA (Life Technologies, Burlington, ON, Canada), and quantified using a Synergy 2 Multimode Microplate Reader (BioTek, Winooski, VT, USA). Total RNA was reverse transcribed using the iScript^TM^ Reverse Transcription Supermix (Bio-Rad, Mississauga, ON, Canada), and diluted five-fold post-synthesis of cDNA with molecular-grade nuclease-free water (dH_2_0) (Wisent Corporation, St. Bruno, QC, Canada). GPA2 and GPB5 transcript abundance was quantified using PowerUp^TM^ SYBR^®^ Green Master Mix (Applied Biosystems, Carlsbad, CA, USA) and analyzed on a StepOnePlus^TM^ Real-Time PCR System (Applied Biosystems, Carlsbad, CA, USA) as described previously (1). RT-qPCR conditions were described previously (2) and GPA2- and GPB5-specific primer sequences were designed in an earlier study (3). GPA2 and GPB5 transcript abundance was normalized to the expression of ribosomal protein 49 (GenBank accession: AY539746) and 60S ribosomal protein S18 (GenBank accession: XM_001660270) following the ΔΔCt method as described previously (3). Measurements were performed using three technical replicates per sample and three biological replicates for each tissue/organ sample.

**Preparation of plasmid constructs for heterologous expression**

Tags were incorporated onto the carboxyl termini with two consecutive PCR amplifications using plasmid miniprep that contained each ORF as the initial template, and PCR amplification was performed using OneTaq^®^ DNA polymerase (New England Biolabs) and primers listed in Table S2. Conditions for the first PCR amplification were as follows: initial denaturation cycle (94 °C, 30 s) and 35 cycles of denaturation (94 °C, 20 s), annealing for *A. aegypti* GPA2 (involving GPA2 Kozak F1/GPA2 FLAG R1 primers) 10 cycles at 45 °C followed by 25 cycles at 57 °C and *H. sapiens* GPB5 (involving GPB5 Kozak F1/GPB5 FLAG R1 primers) at 64 °C, with each cycle lasting 20 s) and extension (68 °C, 30 s), followed by a final extension (68 °C, 5 min). The subsequent PCR amplification to complete the incorporation of each tag consisted of an initial denaturation (94 °C, 30 s) and 35 cycles of denaturation (94 °C, 20 s), annealing for *A. aegypti* GPA2 (involving GPA2 Kozak F1/GPA2 FLAG R2 primers) 10 cycles at 49 °C and 25 cycles at 58 °C, *H. sapiens* GPB5 (involving GPB5 Kozak F1/GPB5 FLAG R2 primers) 10 cycles at 54 °C and 25 cycles at 60 °C, 20 s) and extension (68 °C, 30 s), followed by a final extension (68 °C, 5 min). Tagged PCR amplicons were cloned into pGEM T Easy vector (Promega, Madison, WI, USA) and confirmed for base accuracy by Sanger sequencing (The Centre for Applied Genomics, Sick Kids Hospital, Toronto, ON, Canada). After sequence verification, plasmid clones were *Not*1 digested (Anza™ cloning system; Life Technologies, Burlington, ON, Canada) for 1 h at 37°C, and His- or FLAG-tagged GPA2 and GPB5 sequences were separated by electrophoresis on a 1% agarose ethidium bromide-stained gel using 1x Tris-Acetate EDTA (TAE) buffer. Samples were then gel-purified and ligated to the mammalian expression vector pcDNA3.1^+^ (Life Technologies, Burlington, ON) following a previously described protocol (2).

**Generation of tethered *A. aegypti* GPA2/GPB5 construct**

All PCR conditions involved miniprep samples of the corresponding templates and Q5 High Fidelity DNA Polymerase (New England Biolabs, Whitby, ON) for amplification, following manufacturer recommended conditions. Overall, a portion of the hexa-histidine tagged glycine-serine repeat linker sequence was incorporated onto the amino terminus of *A. aegypti* GPA2 propeptide sequence, and a *Not*1 restriction enzyme and pcDNA3.1^+^ plasmid DNA sequence overhang was made on the carboxyl terminus (see primer list in Table S2). For the *A. aegypti* GPB5 prepropeptide sequence, a portion of the linker sequence was incorporated onto the carboxyl terminus and an *Eco*R1 restriction enzyme and pcDNA3.1^+^ plasmid DNA sequence overhang was made on the amino terminus (see primer list in Table S2). Modified *A. aegypti* GPA2 and GPB5 sequences were amplified in three consecutive PCR reactions (i – iii). PCR reaction conditions included i) 1 cycle of denaturation (98 °C, 30 s) and 35 cycles of denaturation (98 °C, 10 s), annealing (GPA2: tethered GPA2 F1/R1 primers and 70 °C, GPB5: tethered GPB5 F1/R1 primers and 66 °C, 30 s), and an extension (72 °C, 35 s), followed by a final extension (72 °C, 2 min), ii) 1 cycle of denaturation (98 °C, 30 s) and 35 cycles of denaturation (98 °C, 10 s), annealing (GPA2: tethered GPA2 F2/R2 primers and 61 °C, GPB5: tethered GPB5 F2/R2 primers and 62 °C, 30 s) and extension (72 °C, 35 s), followed by a final extension (72 °C, 2 min), iii) 1 cycle of denaturation (98 °C, 30 s) and 35 cycles of denaturation (98 °C, 10 s), annealing (GPA2: tethered GPA2 F3/R3 primers and 70 °C, GPB5: tethered GPB5 F3/R3 primers and 65 °C, 30 s) and extension (72 °C, 35 s), followed by a final extension (72 °C, 2 min) (see Table S2). To generate the expression cassette template, empty pcDNA3.1^+^ expression vector was double digested with Anza^TM^ *Eco*R1 and Anza^TM^ *Not*1 (Life Technologies, Burlington, ON) restriction enzymes for 1 h at 37°C, migrated on a 1% agarose-TAE ethidium-bromide stained gel and gel-purified. Finally, the three products encompassing modified *A. aegypti* GPA2 propeptide, *A. aegypti* GPB5 prepropeptide and digested pcDNA3.1^+^ plasmid was assembled together using HiFi DNA Assembly Master Mix (New England Biolabs, Whitby, ON) using a DNA molar ratio of 1:2 (vector:insert). After transforming NEB 5-α competent *E. coli* cells with this glycoprotein fusion-encoding construct assembly, bacterial colonies were screened with T7 promoter oligonucleotide and gene-specific primers and verified for base accuracy with Sanger sequencing (The Centre for Applied Genomics, Sick Kids Hospital, Toronto, ON, Canada).

**Table S1:** Primers utilized for RNA probe template generation. GenBank accession numbers: BN001241 (*A. aegypti* GPA2), BN001259 (*A. aegypti* GPB5).

| Oligonucleotide name | Oligonucleotide sequence  (5’ – 3’) | Product size (bp) | | Target and Usage |
| --- | --- | --- | --- | --- |
| T7 promoter | TAATACGACTCACTATAG |  | GPA2 and GPB5 sense and anti-sense RNA probe generation for fluorescence *in situ* hybridisation | |
| *Aedes* GPA2 F | ATGGAGTGGCTACGATTTGC | 363 |  |  |
| *Aedes* GPA2 R | GTGTTACCACTGTAAAAAGGACTGA |  |  |  |
| *Aedes* GPB5 F | ATGATCCTAATCTCGGTATGGA | 486 |  |  |
| *Aedes* GPB5 R | AAATCGAATCCGAATACGCTTAG |  |  |  |

**Table S2:** Primers utilized for synthesis of *A. aegypti* GPA2-FLAG, *H. sapiens* GPB5-His and tethered *A. aegypti* GPA2/GPB5 sequences for heterologous expression in mammalian cell lines. GenBank accession numbers: BN001241 (*A. aegypti* GPA2), BN001259 (*A. aegypti* GPB5) and HF564672.1 (*H. sapiens* GPA2).

| Oligonucleotide name | Oligonucleotide sequence  (5’ – 3’) | Product size (bp) | Target and Usage |
| --- | --- | --- | --- |
| *Aedes* GPA2 kozak F1 | GCCACCATGGAGTGGCTACGATTTGCAACA |  | Sense and antisense primer sets for the incorporation of a FLAG tag on the carboxyl terminus of *A. aegypti* GPA2 |
| *Aedes* GPA2 FLAG R1 | GTTACCACTGTAAAAAGGACGATTACAAGGATGAC | 381 |  |
| *Aedes* GPA2 FLAG R2 | AAGGACGATTACAAGGATGACGACGATAAGTAG | 393 |  |
| *Homo* GPB5 kozak F1 | GCCACCATGAAGCTGGCATTCCTCTTCCTT |  | Sense and antisense primer sets for the incorporation of a hexa-histidine tag on the carboxyl terminus of *H. sapiens* GPB5 |
| *Homo* GPB5 His R1 | GGCCACCACGGAGTGTGAGACC°ATCCATCATCACC | 406 |  |
| *Homo* GPB5-His R2 | TGTGAGACCATCCATCATCACCATCACCATTAG | 417 |  |
| Tethered GPA2 F1 | CGCGAAACGTGGCAGAAACCAG | 309 | Sense and antisense primer sets for the incorporation of (i) a portion of a hexa-histidine tag and glycine-serine repeat linker sequence on the amino terminus and (ii) a *Not*1 restriction enzyme and genomic pcDNA3.1 plasmid sequence overhang on the carboxyl terminus of *A. aegypti* GPA2 |
| Tethered GPA2 R1 | GCTCGTGTTACCACTGTAAAAAGGACTAG |  |  |
| Tethered GPA2 F2 | CATCATCACCATCACCATCGCGAAACGTGGCAGAAACCA | 341 |  |
| Tethered GPA2 R2 | GTTACCACTGTAAAAAGGACTAGGCGGCCGCTCGAGT |  |  |
| Tethered GPA2 F3 | TCGGGGTCGGGGTCGCATCATCACCATCACCATCGCG | 366 |  |
| Tethered GPA2 R3 | GACTAGGCGGCCGCTCGAGTCTAGAGGGCC |  |  |
| Tethered GPB5 F1 | GCCACCATGATCAATCTAATCTCG | 492 | Sense and antisense primer sets for the incorporation of (i) an *Eco*R1 sequence and pcDNA3.1 plasmid sequence overhang on the N terminus and (ii) a portion of a hexa-histidine tag and glycine-serine repeat linker sequence on the carboxyl terminus of *A. aegypti* GPB5 |
| Tethered GPB5 R1 | GGAAATCGAATCCGAATACGCT |  |  |
| Tethered GPB5 F2 | GTGGAATTCGCCACCATGATCAATCTAATC | 519 |  |
| Tethered GPB5 R2 | GGAAATCGAATCCGAATACGCTGGGTCGGGGTCGGGGTCG |  |  |
| Tethered GPB5 F3 | ACTAGTCCAGTGTGGTGGAATTCGCCACCATG | 548 |  |
| Tethered GPB5 R3 | CGCTGGGTCGGGGTCGGGGTCGCATCATCACCATCAC |  |  |

**Table S3.** Normalized scores for predicted G-protein coupling specificity for *A. aegypti* LGR1 and *H. sapiens* TSHR by PRED-COUPLE 2.0 (4). Leucine-rich repeat-containing G protein-coupled receptor 1 (LGR1), thyroid-stimulating hormone receptor (TSHR).

| **G Protein** | ***Aedes aegypti* LGR1** | ***Homo sapiens* TSHR** |
| --- | --- | --- |
| Gs | 0.36 | 0.9 |
| Gi/o | 0.97 | 0.94 |
| Gq/11 | 0.9 | 0.84 |
| G12/13 | 0.92 | 0.92 |
